# Angular momentum regulation may dictate the slip severity in young adults

**DOI:** 10.1101/785808

**Authors:** Mohammad Moein Nazifi, Kurt Beschorner, Pilwon Hur

## Abstract

Falls vastly affect the economy and the society with their high cost, injuries, and mortalities. Slipping is the main trigger for falling. Yet, individuals differ in their ability to recover from slips. Mild slippers can accommodate slips without falling, whereas severe slippers indicate inadequate or slow pre-or post-slip control that make them more prone to fall after a slip. Knowing the discrepancies in different kinematic and kinetic variables in mild and severe slippers helps pinpoint the adverse control responsible for severe slipping and falling. This study examined Center of Mass (COM) height, sagittal angular momentum (*H*), upper body kinematics, and the duration of single/double phase in mild and severe slippers for both normal walking and slipping to identify their differences and possible relationships. Possible causality of such relationships were also studied by observing the time-lead of the deviations. Twenty healthy young adults walked in a long walkway for several trials and were slipped unexpectedly. They were classified into mild and severe slippers based on their slip severity. No inter-group differences were observed in the upper extremity kinematics. It was found that mild and severe slippers do not differ in the studied variables during normal gait; however, they do show significant differences through slipping. Compared to mild slippers, sever slippers lowered their COM height following a slip, presented higher *H*, and shortened their single support phase (*p*-value<0.05 for all). Based on the time-lead observed in *H* over all other variables suggests that angular momentum may be the key variable in controlling slips.

## Introduction

During the year 2015, injuries caused by slips, trips and falls were the second leading cause of fatal injuries in the US, for the second consecutive year (1). Fall related mortality aggravates in elderly, with more than 75% of all fall related deaths happening in persons older than 65 (2). The US economy annually sustains a damage of over $180 billion caused by falls (3). While studies argue that slipping is the main trigger to falling (4–6); preventive measures against slipping should be perused more rigorously. More specifically, studying slipping and other factors that contribute to falling would be a logical first step toward “fall prevention” as the ultimate goal.

Studies have argued that upon slipping, the Central Nervous System (CNS) has to react with appropriate signals to avoid falling and retain balance (7). Obviously, failing to provide proper responses to slip would result in falling. To provide a safer experiment environment to study slips, scientists enforced usage of harness system and have developed different indicators of falling instead of an actual fall. These measures mainly consisted of a load cell average force during falling, percentage of body height drop while slipping, slipping distance, and peak slipping velocity where some of them were reported to predict falls with 90-100% accuracy (8–13). For instance, Lockhart et al. (13) claimed that slippers can be classified into mild and severe slippers by the peak heel speed after slipping. Specifically, severe slips are described as slips in which the peak heel speed exceeds 1.44 m/s and severe slippers are more prone to fall (13). Conversely, mild slips are less dangerous and mild slippers can recover from slips without falling compared to their severe slipper counterparts.

Additionally, prior studies have shown that one’s risk of fall is affected by both pre-slip control (gait control) and post-slip response (slip control) (14–17). In other words, mild slippers possess different control techniques for both walking and slipping compared to severe slippers. Needless to say, identification of such differences in kinematics, dynamics, and control of walking and slipping between mild and severe slippers would facilitate diagnosis of severe slippers who naturally have a higher risk of fall. Consequently, numerous studies have tried to identify discrepancies based on individuals’ fall/recovery outcome and/or slip severity. These studies targeted a wide range of variables to detect differences between fallers and non-fallers (i.e. persons who recover from slips), such as kinematic variables (e.g., foot-floor angles, slipping distances) (14,18,19), kinetic variable (torques) (7,20), and neuromuscular variables (activation onsets) (17,21,22).

While numerous studies tried to find potential associations between slip severity and kinetic and kinematic variables, there are still several critical variables that have not been studied and compared between mild and severe slippers. More importantly, to the best of our knowledge, no study has examined if the found relationships were of causal nature rather than simple associations. For instance, numerous studies have studied the lower extremity kinematics and kinetics and their association to severe slipping (14,20,23–28). However, very limited number of studies have examined the association of the slip severity with upper extremity kinematics during walking and slipping although it has been shown that upper body kinematics play an important role during slip control (29). Also, while several studies have argued that COM height and its stability play a key role in prediction of a slip outcome (10,18), very few studies have compared the COM height based on slip severity to find potential differences. In addition to COM height, angular momentum (denoted by *H* from engineering literature), a quantity representing the movement of rotation of an object, is also known to be of paramount importance in gait. Different studies have tried to examine angular momentum manipulation for human gait (30–34). Nevertheless, no studies have tried to compute and compare *H* between mild and severe slippers.

Specifically, since slips mostly result in a backward fall (22), studying angular momentum in the sagittal plane (backward/forward falls are equivalent a rotation in the sagittal plane) is of our interest. Lastly, the length of single and double support phase of the gait and slipping, as another remarkable and deciding gait parameters (24,35), has never been compared in mild and severe slippers to identify the differences. We argue that a study comparing these variables among individuals with different slip severity may address the gap in our knowledge and find possible associations. Also, since COM height has been used as the main indicator of the falls in slip studies (8,10,18), any variable that show a time-lag in its deviations compared to COM height, will be rule out from having causal relationship with falls while a time-lead over COM height deviations would increase the likelihood of causal nature of that variable to falls.

Using slip severity as a representative of one’s risk of fall, the objective of this study is to i) compute the shoulder and elbow joint angles, the COM height, sagittal angular momentum (*H*), and length of single/double support throughout walking and slipping of mild and severe slippers, ii) compare them to identify significant inter-group differences, and iii) compare the time sequence of the variables that show inter-group differences with COM height to find potential cause of the severe slipping. We hypothesize that these measures would differ between mild and severe slippers, indicating the different motor control in kinematics and kinetics of walking and slip in both mild and severe slippers. Also, we hypothesize that at least one of the variables would deviate sooner than COM height drop (i.e. indicator of falls), indicating a causal relationship to severe slipping, and hence, falling.

## Methods

### Subjects

Twenty healthy young adults age (11 males and 9 females, age mean ± SD: 23.6 ± 2.52) participated in this experiment at University of Pittsburgh. Subjects signed a written consent form before participation and were excluded in case of any gait disorder history/condition. The de-identified data were transferred to Texas A&M University for further analysis. Both the experiment and the data analysis were approved by the Institutional Review Board of both Universities.

### Procedures

Participants were asked to walk in a long pathway at their comfortable speed. They were told that the floor was dry such that they were not anticipating any slips. After two or three walking trials, a slippery contaminant (75% glycerol, 25% water) was applied to the walkway to generate and collect a slip trial data (Fig 1). Subjects were told to look away from the walkway after each trial and keep the provided headphones on to minimize the possible contamination noise and hence, the slip expectation. Subjects donned an overhead harness for their safety throughout the trials. Matching size PVC-soled shoes were provided for all participants. During the first few walking trial, the relative location of starting point to the upcoming contamination point was adjusted in a way to have subjects step on the slippery surface with their leading leg during the slipping trial.

**Figure 1:**
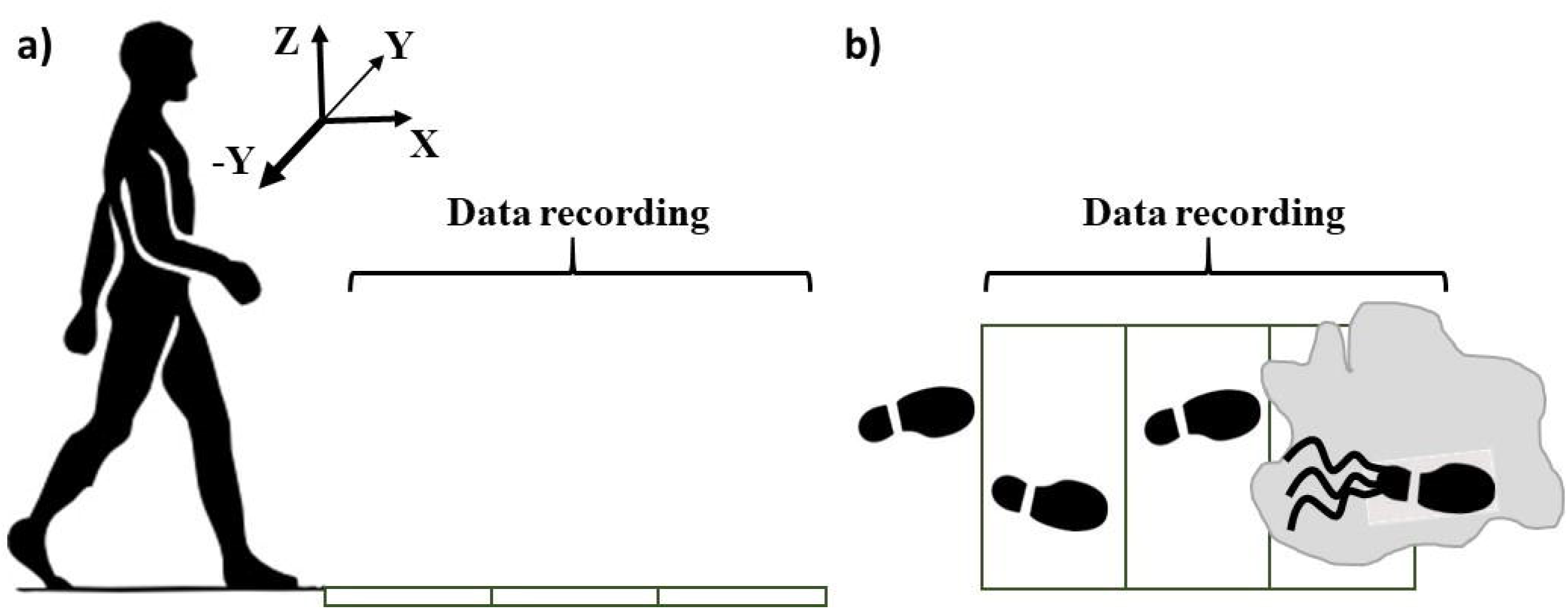
Experimental setup, contamination, and foot placing during the experiment.

### Data and Data Analysis

A set of 79 reflective markers was placed on anatomic bony body landmarks (14) to collect the kinematics at 120 Hz (Vicon 512, Oxford, UK). Subjects’ weight and height were recorded. The markers’ data were low-passed filtered (at 10 Hz) with a second order Butterworth filter (MATLAB, MathWorks, Natick, MA) (20). Using the heel marker information after the slip trial, subjects were classified to mild and severe slippers based on their Peak Heel Speed (PHS) (13,15) to investigate their inter-group differences. Next, based on heel and toe markers, the heel strike and toe-off were calculated and the corresponding double/single support phase of the gait were measured for each individual. The filtered markers data were also used in a generic code (MATLAB, R2017a MathWorks, Natick, MA) to compute limb and joint positions (for both upper and lower extremity) on both right/leading/slipping side (L) and left/trailing/non-slipping side (T). The rotations of the upper extremity joints, the head kinematics, and the hands’ kinematics were not studied as they have little to no effect on the angular momentum. Using anthropometric relative joint and COM positions (36), the center of mass of each limb was calculated and used to measure the position and velocity of the whole body’s center of mass. The center of mass was then normalized using subjects’ heights and presented as a height percentage. Finally, using the same segmental analysis method as COM, the angular momentum of the body was calculated by multiplying the relative velocity of each limb compared to COM to its relative distance to COM and its mass as described in Equation 1:

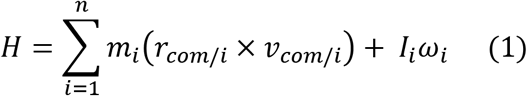

where *m*_*i*_ is the mass of the *i-*th limb, and *r*_*COM/i*_ and ν_*COM/i*_ are the relative distance and velocity of the *i-*th limb with respect to the whole-body COM and *I*_*i*_ and ω_*i*_ are the mass moment of inertia and absolute sagittal plane angular velocity, respectively. According to our reference frame (Fig 1a), a positive angular momentum indicates a general backward rotation whereas a negative *H* shows a forward rotation (30,37). Moreover, *H* is a function of COM velocity (m/s), relative distance of each limb to whole-body-COM (m, function of participant’s height), and mass (kg). Hence, a unitless/non-dimensional *H* was created by dividing original *H* to one’s average COM velocity, mass, and height (31). This would remove subjective differences and make unitless *H* a more appropriate candidate to present inter-subject differences.

Furthermore, to eliminate effect of different gait speeds, gait cycle was normalized to 100 points for each subject to facilitate a point-to-point inter-subject comparison. The comparison was made between a full gait cycle (0% to 100%) for normal *walking* and an additional 30% of gait cycle through *slipping* (100% + 30% = 130% of gait cycle time). 30% of gait cycle time is enough to capture the slip response of the subjects according to (38). Considering the slip to happen at time = 0%, the prior full gait cycle would have happened from −100% to 0%. Also, the slipping would happen starting from 0% and the analysis continued until 30% (Fig 2). The upper body kinematics, the *z* component of the COM (COM height), and the *y* component of *H* (angular momentum in sagittal plane) (Fig 1) were used for comparison between the mild and severe slippers at each percentage of the gait and slipping (i.e. 130 data points). The single/double stance was studied for more than 30% through slipping. Since double stance happens later in a gait cycle, we studied this variable for a full gait cycle before slip initiation (i.e. from −100% to 0%) and a full gait cycle time length after slip initiation (i.e. from 0% to 100%, total of 200% instead of 130%). The data were checked for normality and homogeneity of variance (using Shapiro Wilk and Levene’s test, respectively). Statistical Parametric Mapping (SPM) at significance of 0.05 was used (MATLAB, MathWorks, Natick, MA) to identify the regions of the gait cycle where the upper body kinematics, *H*, and COM height deviate significantly between groups. SPM is a statistical technique that can be used to examine differences observed in time-series data while adjusting for the multiple comparisons (39,40). Moreover, an independent *t*-tests were used to detect statistically significant differences in the single/double stance duration between mild and severe slippers at a significance of 0.05 (SPSS v21, IBM, Chicago, IL) as this variable is not considered a time-series and only presents the time of the transition from single to double stance (The variances were also checked and in case of significant difference in variance, a Welch *t*-test was used instead of an independent *t*-test).

**Figure 2:**
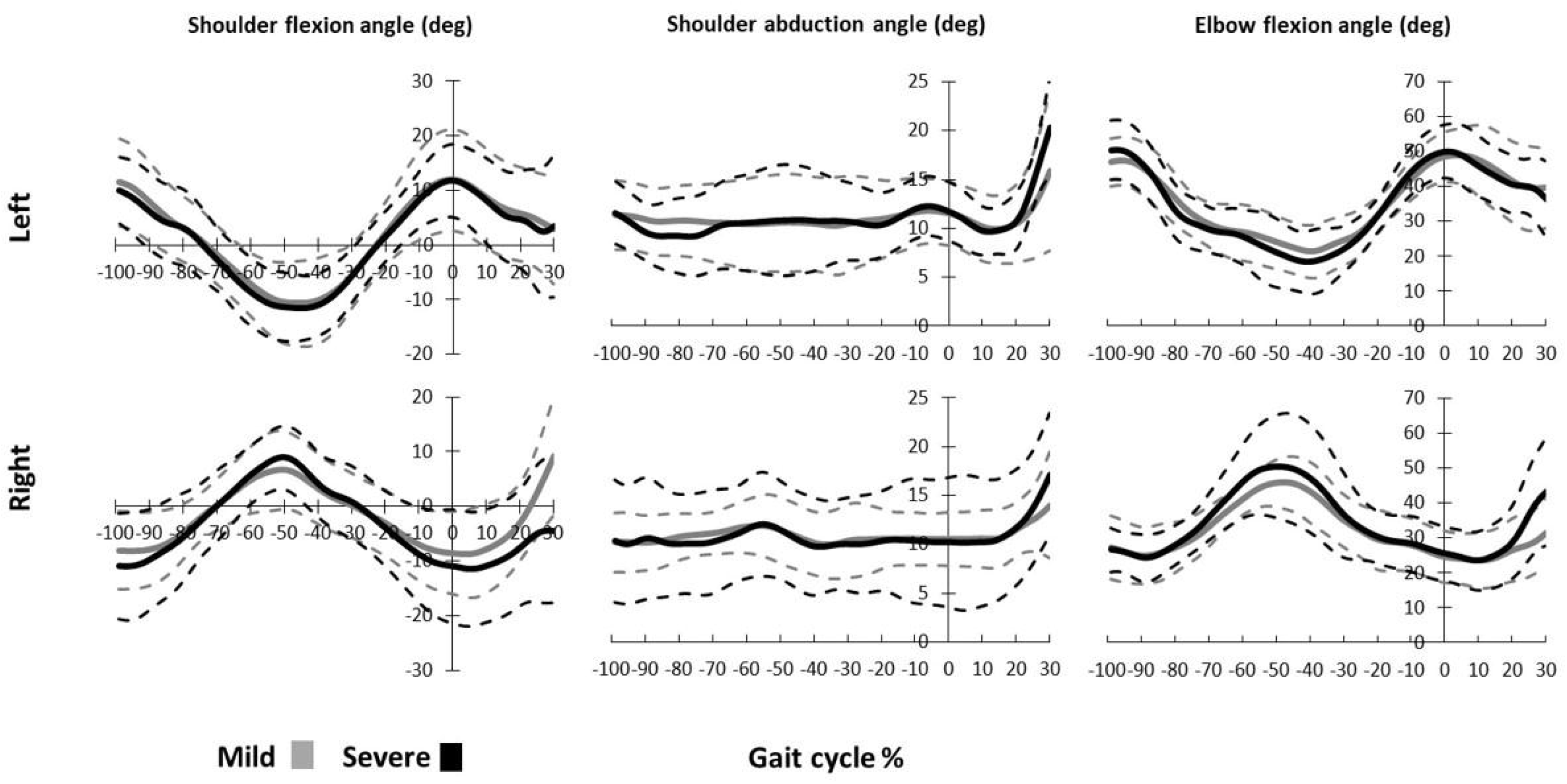
Upper body kinematics for mild and severe slippers for a full gait cycle prior to slip (−100% to 0%) and 30% of the gait cycle time length during slipping.

## Results

Eight of the twenty participants were found to be severe slippers due to their PHS, while the rest were mild slippers. Statistical tests showed no gender, age, or sex related association for slip severity (Table. 1). The upper body kinematics were extracted (Fig 2), and the statistical comparison indicated that there were no significant inter-group differences in the upper body kinematics both before and after the slip initiation, meaning that mild and severe slippers exhibited the same kinematics in their upper extremity before and after slipping.

**Table 1:** Different severity groups’ information. Please note that there was no significant difference in any of the variables at level of 0.05, except PHS.

| Mean±SD | PHS<br>(m/s) | Age | Mass<br>(Kg) | Height<br>(cm) | Sex<br>(M/F) |
| --- | --- | --- | --- | --- | --- |
| Mild | 0.63±0.25 | 24.17±2.79 | 68.41±11.89 | 171.75±8.59 | 5/7 |
| Severe | 1.87±0.27 | 22.75±1.48 | 70.00±11.37 | 175.19±7.57 | 6/2 |

The SPM analysis indicated that mild and severe slippers differ in their COM height and dimensionless sagittal angular momentum after slip initiation. The independent *t*-tests showed that the duration of single/double support differs for different severity groups following slip initiation. Preceding the heel contact on slippery contaminant (i.e. walking), the mild and severe slippers did not differ in COM height; however, from 24%-30% of the gait cycle into slipping, COM height became significantly higher in mild slippers (*p*-value = 0.049) (Fig 3a).

**Figure 3:**
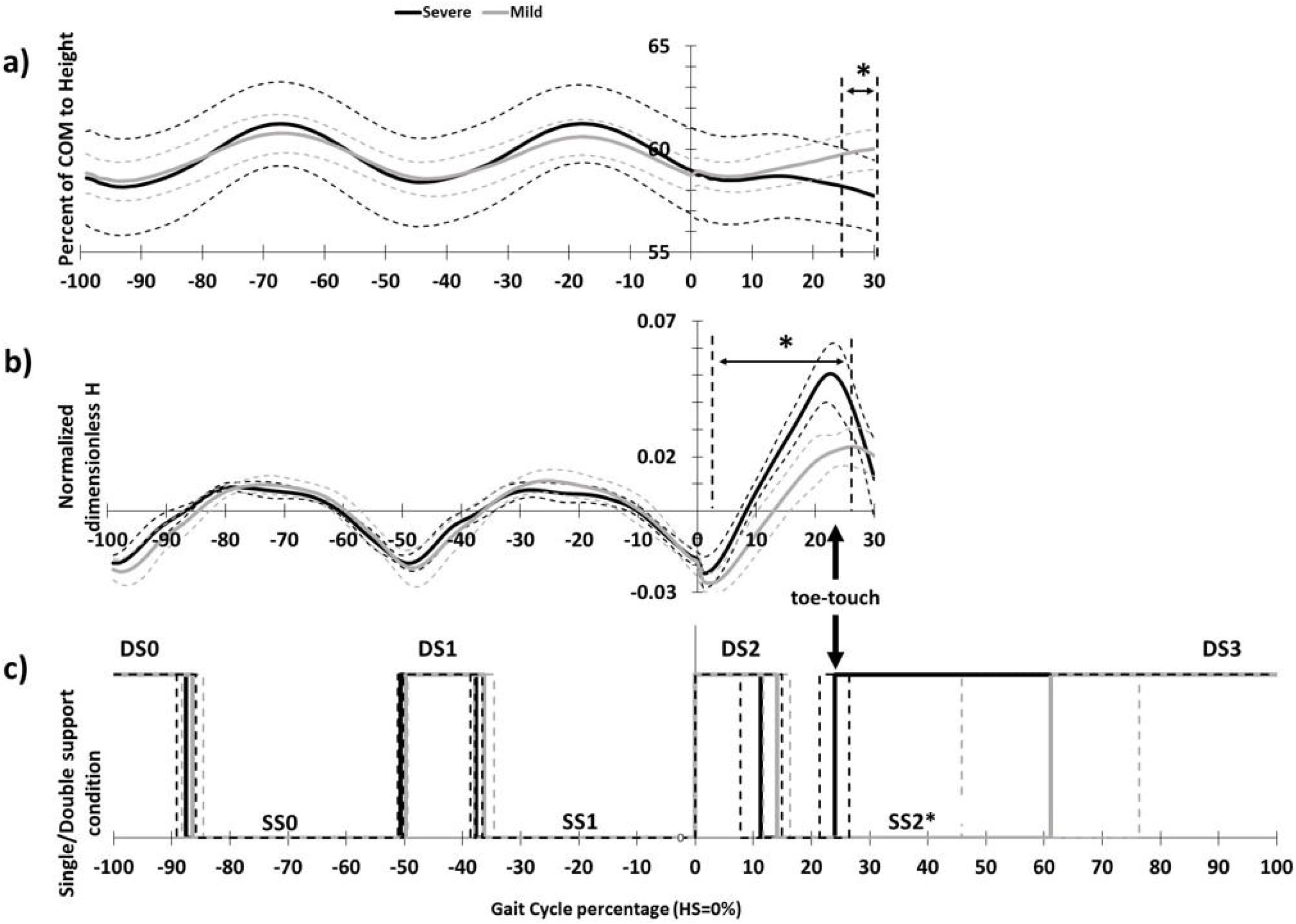
COM height, sagittal *H*, and single/double support phase duration for mild and severe slippers. Asterisks indicate significant difference.

Moreover, for the dimensionless sagittal angular momentum, mild and severe slippers showed a significant difference from 4%-26% into slipping (*p*-value<0.001) (Fig 3b). Lastly, statistical analysis indicated that severe slippers have a shortened single stance phase compared to their mild slipper counter parts, post slip initiation (*p*-value<0.001) (Fig 3c, SS2).

## Discussion

The significant discrepancies in COM height post-slipping, could be interpreted as a strong correlation between slip severity and deviation of COM height. COM height was significantly dropped compared to normal gait following a slip in severe slippers, while mild slippers maintained their post-slip COM height fairly similar to COM height during normal walking (Fig 3a). The observed pattern in mild slippers indicate that subjects who could maintain their post-slip COM height similar to their normal walking COM height, are less likely to experience a severe slip. On the other hand, a sudden decrease in the COM height was associated with severe slipping and hence, falling. Hence, controlling COM could be a useful yardstick in identification of people with high risk of fall due to the identified association and may result in development of rehabilitative/preventative anti-fall devices. This finding stays consistent with previous articles that claimed the height drop can be used as an indicator of falls in presence of harness (8). However, another possible interpretation for the observed deviation between pre-slip and post-slip COM height in severe slippers can be a potential safety strategy. In other words, it is possible that due to the severe slip, the CNS changes its strategy from maintain the COM height to deliberately lowering the COM in order to take a safer fall. This interpretation however, requires further investigation and will be remain unanswered to be studied in our future studies.

Furthermore, the severe slippers experienced a shortened single stance phase following a slip. “Toe-touch” response is a known way to increase the base of support during slipping (15,29). However, it seems that this strategy is only used in more severe slips, since all mild slippers avoided using this strategy while slipping and continued countering slip on one limb, without a toe-touch. Considering this strong association, it is likely that only severe slips required this response to maintain their balance. A more focused study is required to examine this hypothesis and to see if a toe-touch response has a higher trigger for its activation, using an accelerating treadmill that could induce slips with desired intensities.

Analysis on the sagittal angular momentum showed that mild and severe slippers differ in their *H* early after onset of the slip at 4% until 26% of slipping (*p*-value<0.001, Fig 3b). As mentioned before, *H* can be interpreted as a representative of body’s movement of rotation. Human gait exhibits a periodic angular momentum pattern (Fig 3b) and the gait pattern has evolved in a way to matches dynamics of the body while walking and to restrain the *H* by countering the upper body movements (i.e. moving limbs in opposite directions) and usage of the impact of heel strikes (30,31). Modulating the *H* values throughout walking is of crucial importance (30,31). According to our findings, it seems that severe slippers could not modulate and counter their excessive body rotation caused by slipping from 4%-26% into slipping. On the other hand, mild slippers have been able to maintain their angular momentum significantly lower (and more similar to normal walking), which made them more successful in maintaining their balance following a slip.

Association of an excessive *H* with severe slipping and falling suggests that falling does not happen as a vertical COM drop, but it happens as a backward rotational move resulting in a significant vertical COM drop. More importantly, the deviation observed in *H* values (onset at 4% into slipping, Fig 3b) had a significant time-lead over the significant drop observed in COM height (at 24%, Fig 3a). As mentioned before, COM height drop has been introduced as one of the main indicators of falls (8,10,41). Since the deviations in *H* happen before the main indicator of falling (i.e. COM height), we suspect the angular momentum of body to be an earlier indicator of falls and one of the key variables in controlling slips. This finding matches with the existing literature that showed a higher hip flexion angle and knee extension angle to be associated with more severe slips (25) as both contribute to a higher backward angular momentum and hence, a potential backward falling. In postural balance studies, it has been shown that the CNS has the potential to choose different control strategies and employ them for situations with different intensities (i.e. ankle strategy, hip strategy, stepping strategy (42)). Hence, one may speculate that the CNS would react differently to slips with different severities as well (25).

We argue that angular momentum can potentially be a deciding variable in post-slip control, meaning that the CNS may choose different control methods based on *H* value. This hypothesis is substantiated by the pattern observed in the single/double support phase duration. As mentioned, only the severe slippers utilized a ‘toe-touch’ response to their slips. This ‘toe-touch’ response (completed at 23%, Fig 3c) could not have been triggered by COM height drop due to its time-lead (onset at 24%, Fig 3a). Hence, we suggest that this toe-touch response is a measure enforced by the CNS to constrain and regulate the excessive *H* because angular momentum can only be changed by the exertion of an external moment around the body’s COM (which is done by the toe-touch). This is clearly observable in Fig 3b-c, where the excessive positive *H* values in severe slippers (i.e. backward falling) dropped significantly following their toe-touch response that widens base of support to provide moment to prevent backward falling. Further validation of our theory about *H* being a controlling variable in slip control will be an open question for examination for our future studies. Also, we are interested in investigating the angular momentum in other planes in our future studies to further substantiate the current findings.

The upper extremity kinematics stayed consistent with the previous kinematic studies. An arm elevation strategy, as described by (23) was deployed by all subjects (i.e. Fig 2, shoulder abduction happening from 0% to 30%) in response to a slip. This strategy helps moving the COM forward to prevent backward falls, hence subjects tend to move their arms to a more anterior and superior position (i.e. shoulder abduction and flexion, Fig 2, from 0% to 30%) to avoid falls (43,44). However, there were no discrepancies detected between the upper body kinematics for different severities. This indicates that the upper extremity kinematics and control during normal walking and early slipping (up to 30% of the cycle) has little to no significant effect on the slip severity outcome, while many other studies have shown that the lower body kinematics during walking and slipping have a strong association with the slip severity and its outcome (14). Nonetheless, considering our theory of importance of *H*, we suspect the rapid, countermovement of the hands to be a measure to lower whole-body angular momentum. This fact and the timing of this drop in *H* stays consistent with existing literature that suggest upper extremity movements as strategy to prevent falling (29,45).

## Conclusion

This study examined several kinematic and dynamic measures in mild and severe slippers to identify the inter-group differences. We found that mild and severe slippers differ in their control of COM height, sagittal angular momentum, and duration of single/double support phase mainly after slip initiation. Also, the time sequence of the deviations substantiated angular momentum to be a deciding variable in controlling slips. These findings can substantiate that healthy young mild and severe slippers have no difference in their pre-slip control and the higher severity is potentially caused by their post-slip response and probably their angular momentum regulation. Such studies are useful in identification of the underlying causes of severe slipping, which is a main step in fall prevention. Further studies are required to examine these variables in older adults to possibly generalize the findings of this study.

## Acknowledgement

We thank Dr. Rakie Cham for providing the experimental data for the analyses.

